# A multi-sensor system provides spatiotemporal oxygen regulation of gene expression in a *Rhizobium*-legume symbiosis

**DOI:** 10.1101/2020.09.09.289140

**Authors:** Paul J. Rutten, Harrison Steel, Graham A. Hood, Lucie McMurtry, Barney Geddes, Antonis Papachristodoulou, Philip S. Poole

## Abstract

Regulation by oxygen (O_2_) in rhizobia is essential for their symbioses with plants and involves multiple O_2_ sensing proteins. Three sensors exist in the pea microsymbiont *Rhizobium leguminosarum* Rlv3841: hFixL, FnrN and NifA. At low O_2_ concentrations (1%) hFixL signals via FxkR to induce expression of the FixK transcription factor, which activates transcription of downstream genes. These include *fixNOQP*, encoding the high-affinity cbb_3_-type terminal oxidase used in symbiosis. *In vitro*, the Rlv3841 hFixL-FxkR-FixK cascade was active at 1% O_2_, and confocal microscopy showed the cascade is active in the earliest stages of Rlv3841 differentiation in nodules (zones I-II). *In vitro* and *in vivo* work showed that the hFixL-FxkR-FixK cascade also induces transcription of *fnrN* at 1% O_2_ and in the earliest stages of Rlv3841 differentiation in nodules. We confirmed past findings suggesting a role for FnrN in *fixNOQP* expression. However, unlike hFixL-FxkR-FixK, Rlv3841 FnrN was only active in the near-anaerobic zones III-IV of pea nodules. Quantification of *fixNOQP* expression in nodules showed this was driven primarily by FnrN, with minimal direct hFixL-FxkR-FixK induction. Thus, FnrN is key for full symbiotic expression of *fixNOQP*. Without FnrN, nitrogen fixation was reduced by 85% in Rlv3841, while eliminating hFixL only reduced fixation by 25%. The hFixL-FxkR-FixK system effectively primes the O_2_ response by increasing *fnrN* expression in early differentiation (zones I-II). In Zone III of mature nodules, the near-anaerobic conditions activate FnrN, which induces *fixNOQP* transcription to the level required to achieve wild-type nitrogen fixation activity. Modelling and transcriptional analysis indicates that the different O_2_ sensitivities of hFixL and FnrN lead to a nuanced spatiotemporal pattern of gene regulation in different nodule zones in response to changing O_2_ concentration. Multi-sensor O_2_ regulation systems are prevalent in rhizobia, suggesting the fine-tuned control they enable is common and maximizes the effectiveness of the symbioses.

**Author Summary:** Rhizobia are soil bacteria that form a symbiosis with legume plants. In exchange for shelter from the plant, rhizobia provide nitrogen fertilizer, produced by nitrogen fixation. Fixation is catalysed by the nitrogenase enzyme, which is inactivated by oxygen. To prevent this, plants house rhizobia in root nodules, which create a low oxygen environment. However, rhizobia need oxygen, and must adapt to survive low oxygen in the nodule. Key to this is regulating their genes based on oxygen concentration. We studied one *Rhizobium* species which uses three different protein sensors of oxygen, each turning on at a different oxygen concentration. As the bacteria get deeper inside the plant nodule and the oxygen concentration drops, each sensor switches on in turn. Our results also show that the first sensor to turn on, hFixL, primes the second sensor, FnrN. This prepares the rhizobia for the core region of the nodule where oxygen concentration is lowest and most nitrogen fixation takes place. If both sensors are removed, the bacteria cannot fix nitrogen. Many rhizobia have several oxygen sensing proteins, so using multiple sensors is likely a common strategy that makes it possible for rhizobia to adapt to low oxygen gradually in stages during symbiosis.

## 1. Introduction

Rhizobia are alpha-proteobacteria that engage in symbiosis with legume plants [1]. The bacteria convert inert atmospheric N_2_ into biologically accessible ammonia and provide it to their plant host in a process called nitrogen fixation [2,3]. All biological fixation is catalysed by the nitrogenase enzyme complex that evolved before the Great Oxygenation Event and requires near-anoxic conditions to function [4–6]. However, rhizobia are obligate aerobes and must respire to meet the high energy demands of nitrogen fixation [7,8]. These competing requirements create an ‘oxygen paradox’ in symbiotic nitrogen fixation [9,10]. To overcome this paradox, intricate cooperation between rhizobia and their plant partners has evolved (reviewed in [11,12]). Legume plants host rhizobia in dedicated root nodules which have an internal low oxygen (O_2_) environment suitable for nitrogenase activity [13–15]. To achieve this, O_2_ is captured and shuttled to bacteroids by plant leghaemoglobins [16–19]. The concentration of remaining free O_2_ in the core nitrogen fixation zone of nodules is as low as 20-50 nM [20,21]. Inside nodules rhizobia differentiate into quasi-organelle bacteroids specialized for nitrogen fixation [22,23]. Rhizobial regulatory mechanisms that are sensitive to O_2_ tension are essential for successful differentiation into bacteroids and the establishment of a productive symbiosis [24–26].

Multiple O_2_ sensing systems have evolved in rhizobia, three of which are widespread and often co-exist within the same organism [27]. The first system relies on the membrane-bound FixL oxygen sensor which forms a two-component system (TCS) with the FixJ receiver protein (reviewed in [28,29]). Under microaerobic conditions, FixL phosphorylates FixJ, which in turn induces expression of the gene for the FixK transcription factor [30–32]. FixK induces expression of downstream genes by binding as a dimer to an ‘anaerobox’ motif (TTGAT-N_4_-ATCAA) upstream of their promoters [33,34].

The second common O_2_ regulation system is based on a variant of FixL called hybrid FixL (hFixL) [35,36]. This forms an alternative TCS with FxkR acting as the receiver protein. FxkR is not a FixJ homolog but similarly induces expression of *fixK*, by binding to an upstream ‘K-box’ motif (GTTACA-N_4_-GTTACA) [37]. The third O_2_ sensing system is based on the FnrN transcription factor. Like FixK, FnrN binds the anaerobox motif as a dimer and both are close homologs of the *E. coli* anaerobiosis regulator FNR [38–40]. Unlike FixK but like FNR, FnrN contains an N-terminal cysteine-rich cluster that makes the protein a direct sensor of O_2_ [41–44]. The FixL and hFixL sensors are known to become active at relatively mildly microaerobic conditions, including in free-living rhizobia [45–47]. FnrN is likely to be far less O_2_ tolerant. The O_2_ sensitivity of FnrN has not been determined, but the *E. coli* FNR homolog is active only under anaerobic conditions [48–50]. All symbiotic rhizobia studied to date employ at least one of these three systems [11]. It is common for these systems to coexist, notably in *Rhizobium leguminosarum* biovar *viciae* VF39, multiple strains of *Ensifer meliloti* (previously *Sinorhizobium meliloti)* and *Rhizobium etli* CFN42 [35,36,51–53].

The significance of these multi-sensor arrangements, and the relationship between coexisting O_2_ regulators, remains poorly understood. There appears to be a spectrum among rhizobia with some arranging their sensors in ‘fragile’ hierarchical systems where one or more regulators is key, and others having redundant parallel systems in which multiple regulators apparently carry out the same function [54,55]. In *R. leguminosarum* bv. VF39, knocking out FnrN or hFixL reduced nitrogen fixation to 30% or 50% of WT respectively, suggesting a non-hierarchical, redundant arrangement [51]. By contrast *R. etli* CFN42 appears to employ a complex hierarchical system, with multiple homologs of FixK and FnrN regulating each other’s expression [52,53]. The hFixL system of *R. etli* CFN42 is dispensable as a double *fixK* mutant had no effect on nitrogen fixation, whilst a double *fnrN* mutant reduced fixation to 20% of WT levels. Species encoding homologs of only hFixL or FnrN have also been found. *Rhizobium leguminosarum* biovar *viciae* UPM791 contains two FnrN homologs but neither FixL nor hFixL [56]. It is unknown whether the two FnrN proteins respond to different O_2_ concentrations or act in a redundant fashion. *E. meliloti* 1021 contains no FnrN homolog but a well-studied FixLJ system and appears to have homologs of hFixL and FxkR [57,58].

To examine the relationship between hFixL and FnrN, we studied the model organism *Rhizobium leguminosarum* biovar *viciae* 3841 (Rlv3841) which employs both systems (Figure 1) [59,60]. Rlv3841 encodes two copies of hFixL, which we named hFixL_9_ (pRL90020) and hFixL_c_ (RL1879). The strain also contains two homologs of FxkR, FxkR_9_ (pRL90026) and FxkR_c_ (RL1881). It has three putative FixK proteins, which we designated FixK_9a_ (pRL90019), FixK_9b_ (pRL90025) and FixK_c_ (RL1880). Both *fixK*_9a_-*hfixL*_9_ and *fixK*_c_-*hfixL*_c_ are in putative operons (Figure 1B and 1D). Rlv3841 has one homolog of FnrN, regulated by two anaeroboxes. A similar dual-anaerobox arrangement exists in Rlv UPM791, where FnrN positively and negatively auto-regulates its own expression [61]. Binding of FnrN to the distal anaerobox induces *fnrN* transcription and binding to the proximal anaerobox represses it. Auto-activation of *fnrN* has also been reported in *Rhizobium etli* CNPAF512 [62]. FixK regulation of *fnrN* expression is likely as it also binds anaeroboxes, but this had not been investigated. A parallel arrangement of the hFixL-FxkR-FixK and FnrN systems would produce redundancy, whereas an arrangement in series would create hierarchy between the systems. Our goal is therefore to understand how the two O_2_-sensing systems interact in Rlv3841 and to provide insight into why they coexist.

**Fig. 1.**
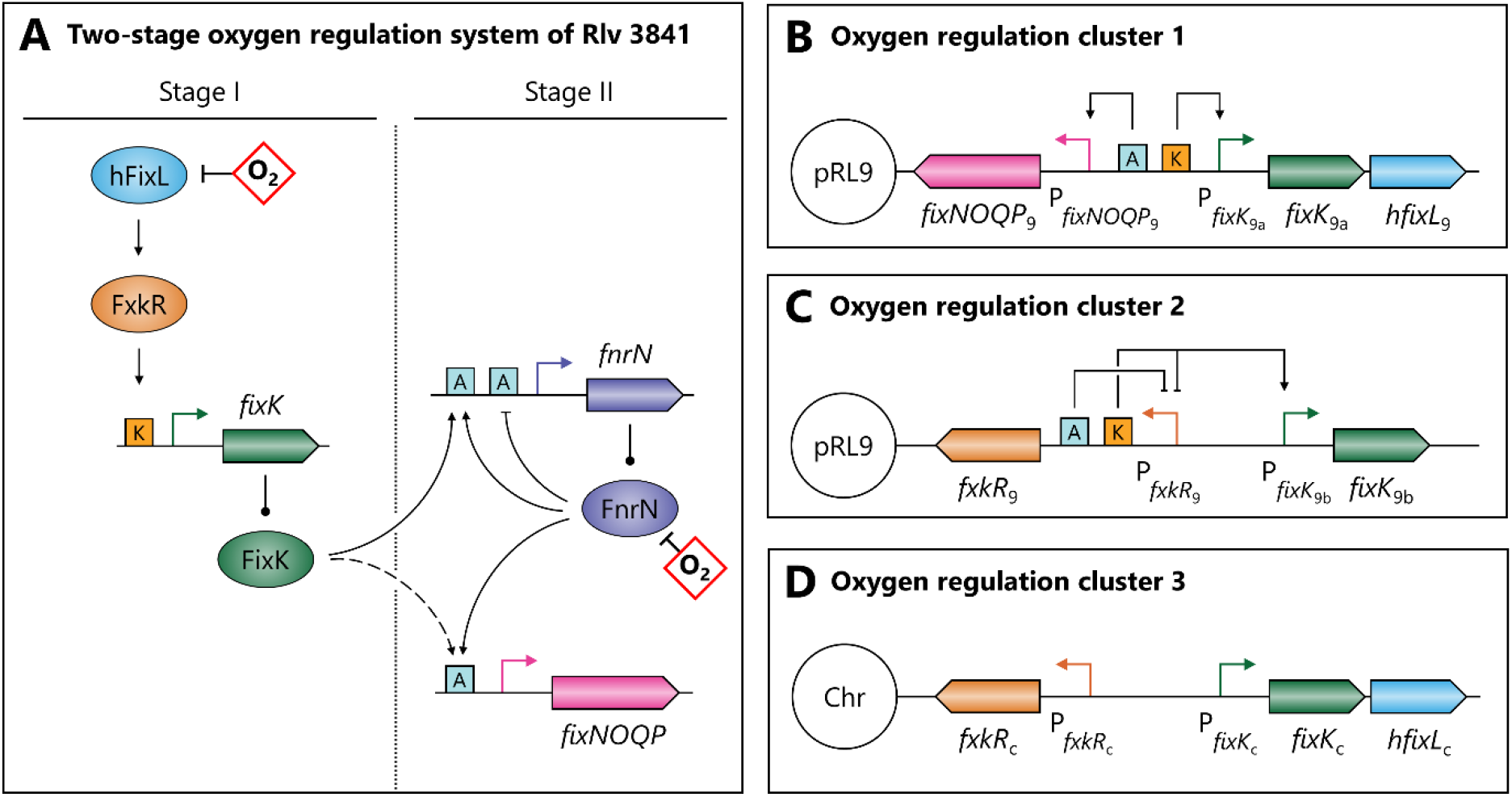
The integrated hFixL-FxkR-FixK and FnrN oxygen regulation systems of Rlv3841 form a single pathway and are genetically clustered. Oxygen is shown in red diamonds. Proteins are shown as ovals, operator sites as squares and genes as pointed rectangles. Promoters are shown as right-angled arrows. Line endings indicate activation (arrows), inhibition (blunt end) and translation (circle). **(A)** The single pathway formed by the two systems acts in two stages. Stage I starts under microaerobic conditions and can function outside the nodule. In this stage, hFixL is active but FnrN is not. hFixL activates FxkR, which binds to the K-box operator (orange squares) to induce expression of *fixK*. FixK binds to anaerobox operators (blue squares) to induce expression, including of *fnrN* and *fixNOQP* (dashed line). Once oxygen reaches near-anaerobic conditions inside the nodule, FnrN becomes active and stage II begins. Like FixK, FnrN binds anaeroboxes. It auto-regulates *fnrN* both positively and negatively and induces *fixNOQP* expression. **(B)** Rlv3841 has multiple copies of many oxygen regulation genes and many are arranged in clusters. On plasmid pRL9, *fixK*_9a_ forms an operon with *hfixL*_9_, both regulated by a K-box. This operon is adjacent to *fixNOQP*_9_, regulated by an anaerobox. **(C)** *fixK*_9b_ and *fxkR*_9_ are adjacent, with an anaerobox and a K-box in their intergenic region. Both operators are downstream of the *fxkR*_9_ promoter and likely repress it. The K-box also likely serves to induce *fixK*_9b_ expression. **(D)** The Rlv3841 chromosome also has a cluster, containing *fxkR*_c_, *fixK*_c_ and *hfixL*_c_. Unlike the similar clusters on pRL9, the intergenic region of this cluster contains no anaerobox or K-box operators.

## 2. Results

### 2.1. Expression of *fnrN* is auto-regulated and controlled by the hFixL-FxkR-FixK pathway

The hFixL-FxkR-FixK system is known to be active at relatively high O_2_ concentrations, including in free-living rhizobia under microaerobic conditions [36,63]. The role of FnrN is less well understood and we began by investigating this sensor. The *fnrN* gene contains two anaeroboxes upstream of its promoter, and indeed was induced in response to microaerobic conditions in free-living Rlv3841 (Figure 2). Both FixK and FnrN bind to anaerobox operators and induce gene expression under microaerobic conditions [64,65]. Therefore, microaerobic induction of *fnrN* could be due to FnrN auto-activation and/or induction by the hFixL-FxkR-FixK system. To determine their respective importance, expression of *fnrN* was studied in Rlv3841 mutants defective in either hFixL or FnrN regulation.

**Fig. 2.**
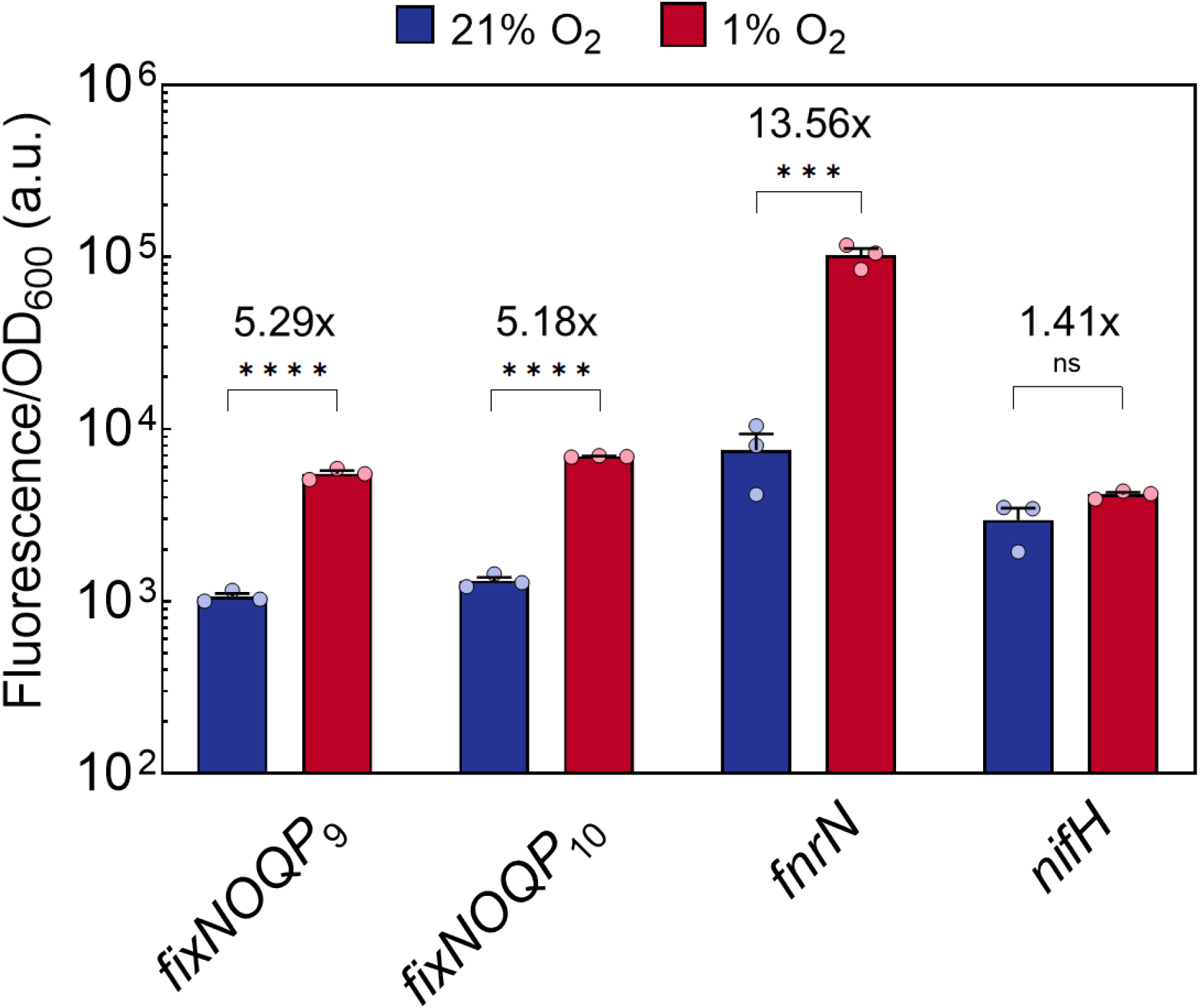
Microaerobiosis induces *fixNOQP* and *fnrN* genes in free-living Rlv3841. Expression of *fixNOQP*_9_, *fixNOQP*_10_ and *fnrN* was increased at 1% O_2_ relative to 21% O_2_. A similar fold induction was recorded for the *fixNOQP* operons, but *fnrN* showed more than double this fold change indicating stronger induction. No effect of O_2_ concentration on *nifH* expression was observed. Values are plotted on a logarithmic scale. Data are averages (±SEM) from three biological replicates, ns (not significant) P ≥ 0.05; ***P < 0.001; ****P < 0.0001; by Student’s t test.

In a double *hfixL* mutant (LMB496; *hfixL*_9_::ΩSpec *hfixL*_c_:pK19 single recombination), free-living microaerobic expression of *fnrN* was reduced to 25% of its wild-type (WT) level (Figure 3). The single mutant of *hfixL*_9_ (LMB495; *hfixL*_9_::ΩSpec) individually reproduced most of this reduction whilst the single mutant of *hfixL*_c_ (LMB403; *hfixL*_c_:pK19 single recombination) did not reduce expression. This suggests hFixL_9_ is the critical hFix_L_ protein under free-living microaerobic conditions, with hFixL_c_ playing little to no role. hFixL acts through the FxkR intermediary in rhizobia and Rlv3841 contains two FxkR homologs [35]. *fxkR*_9_ forms an O_2_ regulation cluster with *fixK*_9b_ (Figure 1C) and *fxkR*_c_ forms a cluster with *fixK*_c_-*hfixL*_c_ (Figure 1D). The first cluster contains an anaerobox and a K-box, but the second cluster contains neither (Figure 1D). Thus, FxkR_9_ is likely the main FxkR protein and *fxkR*_9_ was deleted to produce strain OPS1808 (Δ*fxkR*_9_). This mutant reproduced the reduced induction of *fnrN* under free-living microaerobic conditions observed in the double *hfixL* mutant (Figure 3). This finding supports the role of FxkR_9_ as the mediator of hFixL O_2_ regulation in Rlv3841, in agreement with studies in other rhizobia [35,36]. Studying the role of the hFixL-FxkR-FixK system in *fnrN* expression *in planta*, we observed that the double *hfixL* mutant reduced *fnrN* expression to 28% of WT levels (Figure 4A). This indicates the system also plays an important role in inducing *fnrN* during symbiosis.

**Fig. 3.**
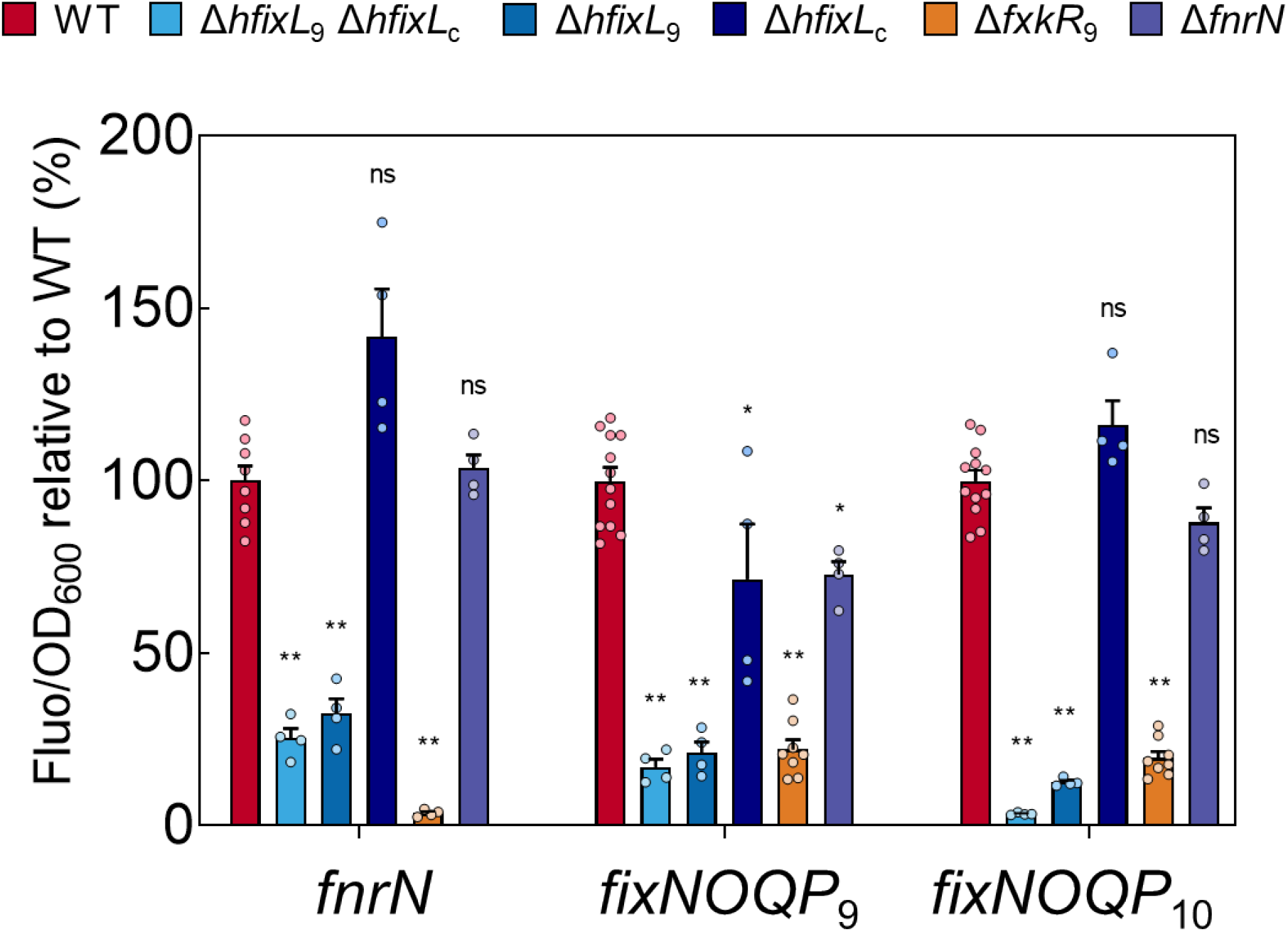
Under *in vitro* microaerobic (1% O_2_) conditions, the hFixL-FxkR-FixK system and not FnrN is a key activator of anaerobox controlled genes. Individual values (Fluo/OD_600_) are normalised such that the WT average is 100% for each reporter. Activity from all three promoters was critically reduced or nearly abolished in the double *hfixL* and *fxkR* mutant backgrounds. The *hfixL*_9_ homolog had a far more pronounced effect on expression of all three genes than did the *hfixL*_c_ homolog. Little or no reduction in expression was observed when *fnrN* was mutated. Data are averages (±SEM) from at least four biological replicates. Statistical tests are differences relative to wild-type (WT) expression; ns (no significant decrease) P ≥ 0.05; *P < 0.001; **P < 0.0001; by one-way ANOVA with Dunnett’s post-hoc test for multiple comparisons.

**Fig. 4.**
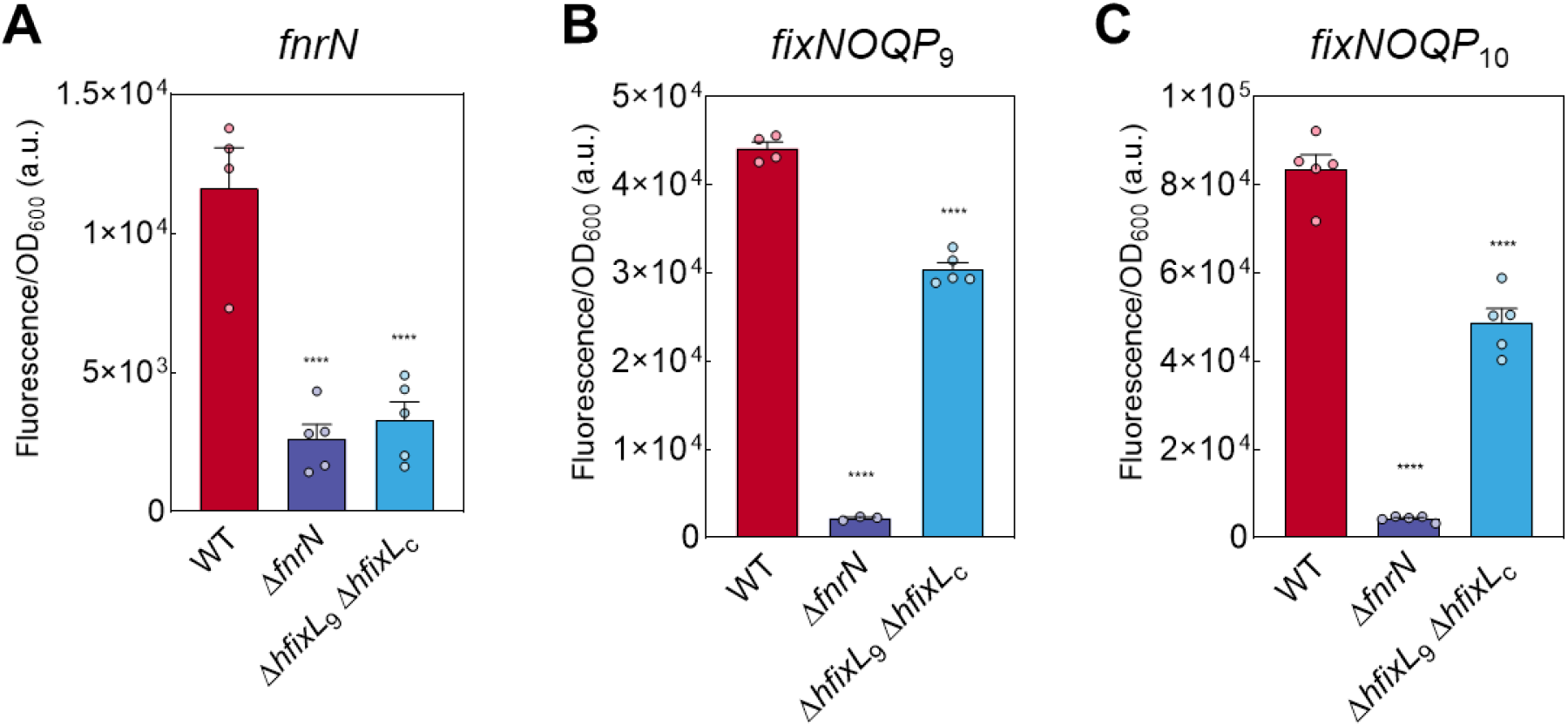
*In planta, fnrN* is both auto-regulated and controlled by the hFixL-FxkR-FixK system, whilst the *fixNOQP* operons are primarily controlled by FnrN. Expression of (**A**) *fnrN* is impaired in the *fnrN* background where auto-activation cannot take place. Expression of *fnrN* is similarly impaired in the double *hfixL* mutant, indicating the hFixL-FxkR-FixK system also plays an important role in symbiotic *fnrN* induction. Expression of (**B**) *fixNOQP*_9_ and (**C**) *fixNOQP*_10_ is significantly reduced in the double *hfixL* mutant and almost abolished in the *fnrN* mutant. Thus both FnrN and the hFixL-FxkR-FixK system play an important role in the expression of all three genes. Data are averages (±SEM) from at least three plants, ****P < 0.0001; by one-way ANOVA with Dunnett’s post-hoc test for multiple comparisons.

We next studied the role of FnrN auto-regulation. A mutant of *fnrN* (LMB648, *fnrN*::ΩTet) had no effect on expression of *fnrN* at 1% O_2_ (Figure 3), indicating FnrN auto-activation does not occur under free-living microaerobic conditions. By contrast, *in planta* the *fnrN* mutant reduced *fnrN* expression to 22% of WT levels (Figure 4A), similar to the reduction observed in the *hfixL* double mutant. FnrN auto-activation is thus an important regulatory effect during symbiosis but not under microaerobic conditions. During symbiosis, expression of *fnrN* is driven by both auto-activation and the hFixL-FxkR-FixK TCS, and both are required to attain full WT expression of the gene.

### 2.2. FnrN is critical for symbiotic gene expression but hFixL also plays an important role

To understand the respective importance of FnrN and hFixL as regulators of anaerobox controlled genes during symbiosis, their role in *fixNOQP* expression was studied. The *fixNOQP* operon encodes a high-affinity *cbb*_3_-type terminal oxidase required for respiration during symbiosis [66–68]. It is typically regulated by an anaerobox in rhizobia, and indeed the operator is present upstream of *fixNOQP* in Rlv3841 [69]. Some rhizobia encode multiple redundant terminal oxidases controlled by different regulator systems, but no alternatives appear to be encoded by Rlv3841 [70,71]. It is therefore likely that the strain relies entirely on *fixNOQP* for respiration during symbiosis. Three putative homologs of *fixNOQP* exist in Rlv3841, but only FixNOQP_9_ (encoded on pRL9) and FixNOQP_10_ (encoded on pRL10) appear functional, as only their genes contain four open reading frames and an upstream anaerobox.

Both *fixNOQP* operons were induced *in vitro* under microaerobic conditions, confirming their regulation by O_2_ (Figure 2). *In vitro* the double *hfixL* mutant severely reduced this microaerobic induction, resulting in minimal expression of *fixNOQP*_10_ and 17% of WT *fixNOQP*_9_ expression (Figure 3). The single *hfixL*_9_ mutant significantly reduced expression of both operons whilst the *hfixL*_c_ mutant only reduced expression of *fixNOQP*_9_, to 71% of WT. These results indicate hFixL_9_ is the dominant protein and hFixL_c_ plays only a minor role in *fixNOQP* expression, matching their respective importance for microaerobic *fnrN* expression (Figure 3). In the Rlv3841 *fxkR*_9_ mutant, expression of both *fixNOQP* operons was reduced to less than 25% of WT, indicating that the protein is required for hFixL regulation of *fixK* and hence *fixNOQP*, as found in other rhizobia [35]. The hFixL-FxkR-FixK system is thus a key regulator of *fixNOQP* expression under free-living microaerobic conditions. By contrast, in these conditions the *fnrN* mutant minimally affected *fixNOQP* expression, with only *fixNOQP*_9_ showing a small albeit statistically significant reduction (73% of WT). In line with our study of *fnrN* expression, the hFixL-FxkR-FixK system is crucial for *fixNOQP* expression under free-living microaerobic conditions whilst FnrN plays a minimal role. It is likely that the FnrN protein remains mostly inactive at the O_2_ concentration (1% O_2_) used in our free-living experiments. Likewise, for comparison we checked the activity of NifA, a central activator of nitrogen fixation genes [72–74]. NifA is O_2_ sensitive and in most rhizobia is active only in the near-anoxic core of nodules [75–79]. We checked the NifA dependant induction of *nifH*, which encodes a component of the nitrogenase complex [80–82]. As expected *nifH* expression did not increase under microaerobic conditions (Figure 2), indicating Rlv3841 NifA is not expressed or inactive under these conditions.

Next, the role of FnrN and hFixL on expression of *fixNOQP in planta* during symbiosis was studied. We found that nodules formed by the *fnrN* mutant expressed both *fixNOQP* operons at only 5% of WT (Figure 4B-C). The FnrN sensor is thus critical for *fixNOQP* expression inside the nodule. In nodules infected by the double *hfixL* mutant, expression of *fixNOQP*_9_ and *fixNOQP*_10_ was reduced to 68% and 58% of WT, respectively. Expression of *fixK*_9a_ was abolished (Supplementary 1, Figure S1), suggesting minimal FixK production in the absence of hFixL-FxkR TCS activity. Taken together, our results indicate that FnrN is critical for *fixNOQP* expression during symbiosis but the hFixL-FxkR system also plays a significant role.

To assess the impact of FnrN and hFixL on symbiotic nitrogen fixation, acetylene reduction assays were performed on pea plants inoculated with O_2_ regulation mutants. In line with its poor expression of *fixNOQP*, the *fnrN* mutant was critically impaired in nitrogen fixation, reducing acetylene at only 15% of the WT level (Figure 5A). Plants inoculated with this mutant produced only small, pale nodules indicative of poor development and low leghaemoglobin production (Figure 5D). Thus, FnrN is critical for effective nitrogen fixation by Rlv3841. Complementation restored 88% of WT acetylene reduction activity and produced nodules indistinguishable from WT (Supplementary 1, Figure S2).

**Fig. 5.**
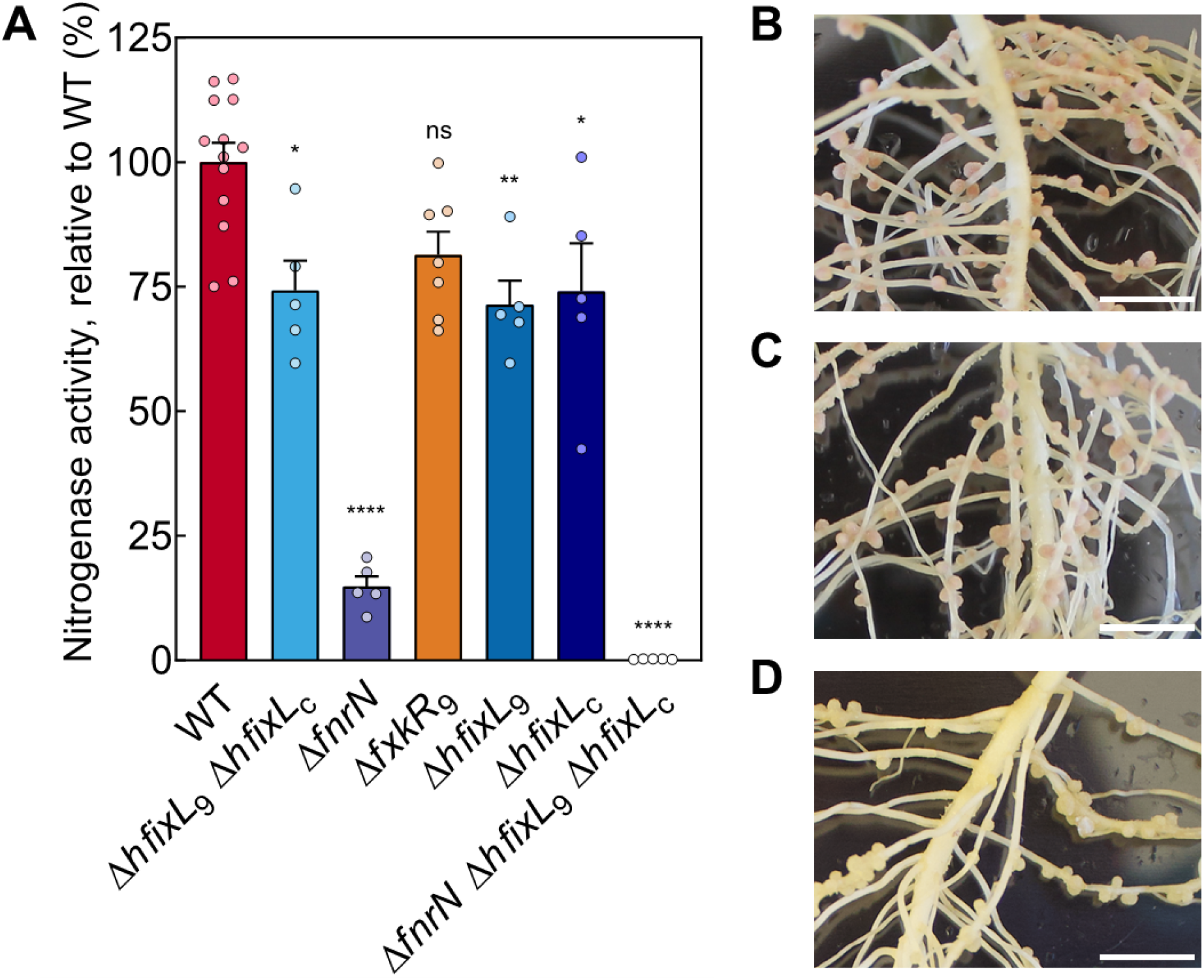
Effect of oxygen regulation mutants on nodule morphology and acetylene reduction rates. **(A)** Acetylene reduction rates of Rlv3841 mutant strains, normalised by WT activity (5.8 μmoles ethylene plant^-1^ hr^-1^, 16.8×10^-1^ μmoles ethylene mg^-1^ of nodules hr^-1^). Knocking out the *hfixL* genes individually and in combination only slightly reduced fixation. The *fnrN* mutant critically reduced fixation. The single *fxkR*_9_ mutant did not significantly reduce fixation (p=0.0584), possibly because of redundancy through the *fxkR*_c_ homolog. The mutant lacking both the FnrN and hFixL-FxkR-FixK systems fixed at only a negligible rate. Rates are normalised per plant to total mass of nodules. Data are averages (±SEM) from at least five plants, ns (not significant) P ≥ 0.05; *P < 0.05; **P < 0.01, by one-way ANOVA with Dunnett’s post-hoc test for multiple comparisons. Photos of nodules colonized by **(B)** WT, **(C)** the double *hfixL* knockout and **(D)** the *fnrN* knockout. Scale bar, 1 cm.

Plants inoculated with either individual *hfixL* mutants or the double mutant were also impaired in nitrogen fixation but retained approximately 75% of WT acetylene reduction activity (Figure 5A). No morphological changes were observed in these nodules (Figure 5C). Thus, the hFixL-FxkR-FixK system is also an important contributor to symbiotic fixation activity and is required to attain a WT level of fixation. Complementation of the double *hfixL* mutant was attempted but the gene was found to be toxic in *E. coli*. The *fxkR*_9_ mutant impaired acetylene reduction rates but the decrease was insufficient to be significant (p = 0.0671). The FxkR_c_ homolog is likely at least partially active and sufficiently produced to rescue hFixL regulation in the absence of FxkR_9_. In the triple *fnrN hfixL*_9_ *hfixL*_c_ mutant (LMB673; *fnrN*::ΩTet *hfixL*_9_::ΩSpec *hfixL*_c_:pK19) only negligible levels of fixation were recorded. This reinforces the importance of the contribution from both systems and suggests no additional regulators exist which induce these anaerobox controlled genes in Rlv3841 during symbiosis.

### 2.3. hFixL and FnrN are active in spatially distinct nodule zones during symbiosis

Legume nodules create a large internal O_2_ gradient, with semi-aerobic conditions at their tip and near-anoxic conditions as low as 20 nM O_2_ in the central nitrogen fixing zone [83,84]. This gradient is typically split into four zones (Figure 6A) containing different O_2_ concentrations and rhizobia in different stages of differentiation (for reviews, see [85,86]). To understand how the hFixL-FxkR-FixK and FnrN systems operate in this context, we used confocal microscopy to map the spatial expression of *fnrN* and *fixNOQP* in nodules.

**Fig. 6.**
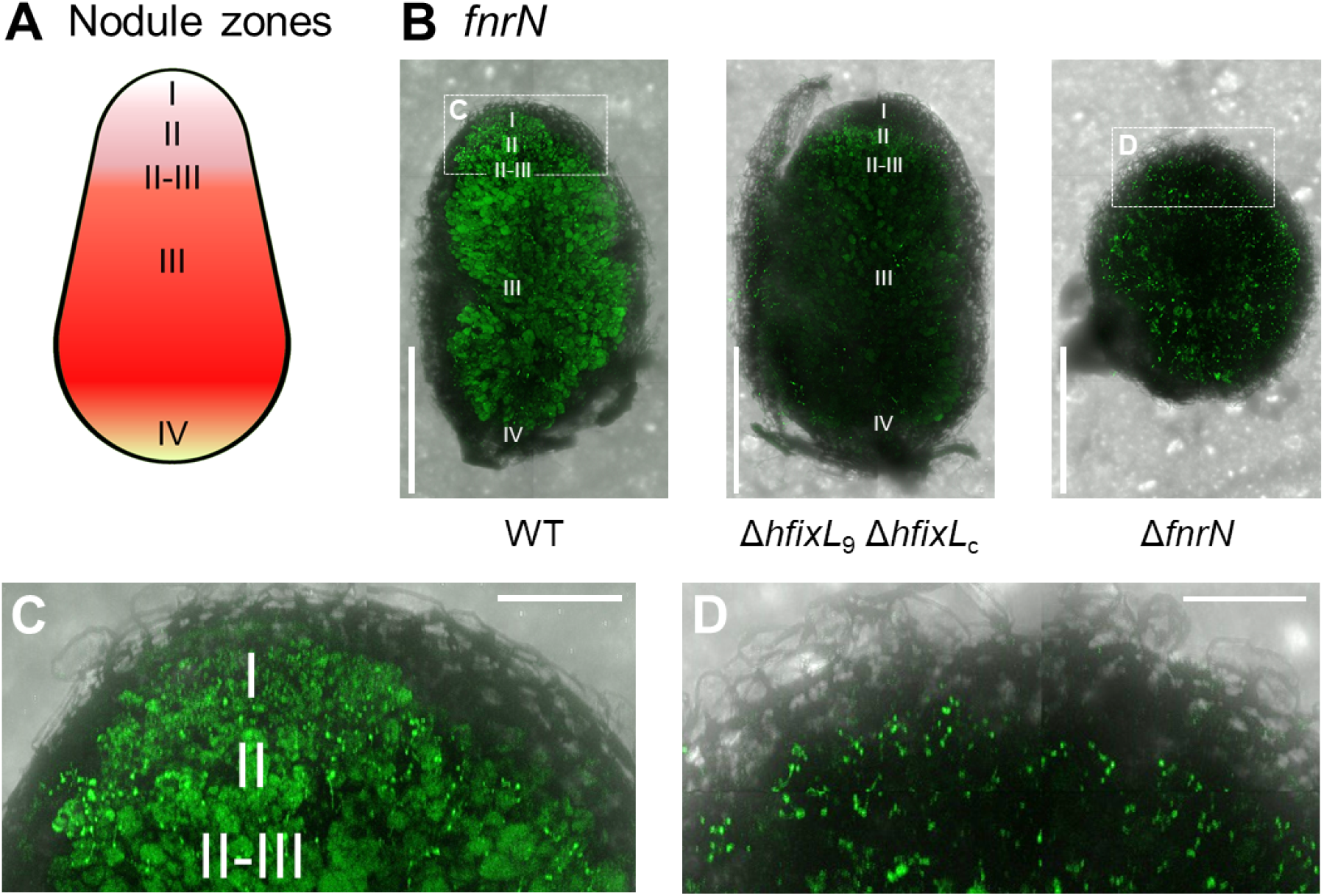
Spatial expression pattern of *fnrN* in nodules infected with Rlv3841 WT and mutants. (**A**) Schematic representation of an indeterminate nodule formed by *P. sativum*. Zone I contains undifferentiated rhizobia in infection threads. Rhizobia enter plant cells in zone II and undergo final differentiation into bacteroids in the II-III interzone. Zone III is the main nitrogen fixing zone. Zone IV contains bacteroids which are beginning to senesce. (**B**) Nodule cross sections showing expression of *fnrN* when inoculated with strains of Rlv3841. Expression begins immediately in zone I in nodules inoculated with WT; see C for a close up of the region highlighted in white. A similar level of expression is present across all zones. When inoculated with the double *hfixL* mutant, expression began in zone II and was highest in this zone. In nodules inoculated with the *fnrN* mutant, expression was observed in infection threads around the periphery of the nodule; see D for a close up of the region highlighted in white. This mutant does not form mature nodules, and the normal zones are therefore unlikely to be fully developed. All three images were captured and processed using identical parameters; see Materials and Methods for details. Scale bar 1 mm. (**C**) Magnified view of *fnrN* expression in WT bacteria in zone I of the nodule. (**D**) Magnified view of *fnrN* expression in the infection threads of a nodule inoculated with the *fnrN* mutant. Scale bar for C and D, 0.25 mm. See Materials and Methods for microscopy technique details.

Expression of *fnrN* in nodules infected with WT Rlv3841 (Figure 6B) was visible throughout all nodule zones. This included expression in infection threads in zone I, indicating that low O_2_ induction of *fnrN* begins when Rlv3841 first enters the nodule and before the bacteria have differentiated into bacteroids. By contrast, *fnrN* expression in zone I was greatly reduced in nodules infected with the double *hfixL* mutant (Figure 6B). This suggests the O_2_ concentration in the relatively aerobic environment of zone I is sufficiently low to activate the hFixL-FxkR-FixK system. In the absence of this system, some *fnrN* expression was retained in zone II and interzone II-III, but this was weaker than WT. Minimal *fnrN* expression was observed in zone III in the *hfixL* double mutant.

In the *fnrN* mutant, expression of *fnrN* appeared to be localized primarily in infection threads, around the entire periphery of the nodule (Figure 6B). Nodules infected by this mutant were severely impaired in their development, failed to elongate and contained little to no leghaemoglobin (Figure 5D). Free O_2_ concentration is unlikely to drop as much in these nodules as it does in fully developed nodules. It is therefore noteworthy that the hFixL-FxkR-FixK system is nevertheless active, suggesting even poorly developed nodules produce a sufficiently low O_2_ concentration to activate it.

Expression of *fixNOQP*_9_ (Figure 7A) and *fixNOQP*_10_ (Figure 7B) followed a similar pattern. In WT Rlv3841, expression of both started abruptly in the II-III interzone of nodules, in agreement with past studies [46,47,87,88]. This abrupt start was absent in nodules infected with the double *hfixL* mutant, indicating it requires the hFixL-FxkR-FixK system. Without hFixL-FxkR-FixK, expression of *fixNOQP*_9_ and *fixNOQP*_10_ started gradually after the II-III interzone, presumably driven by FnrN. Expression was also weaker than the WT. In the *fnrN* mutant, we observed minimal expression of *fixNOQP*_9_. This confirms that the hFixL-FxkR-FixK system can directly induce only minimal *fixNOQP* expression in zone III of mature nodules. Nevertheless, although FnrN is the main driver of *fixNOQP* expression, the hFixL-FxkR-FixK system is required for full *fnrN* expression and therefore also indirectly plays an important role.

**Fig. 7.**
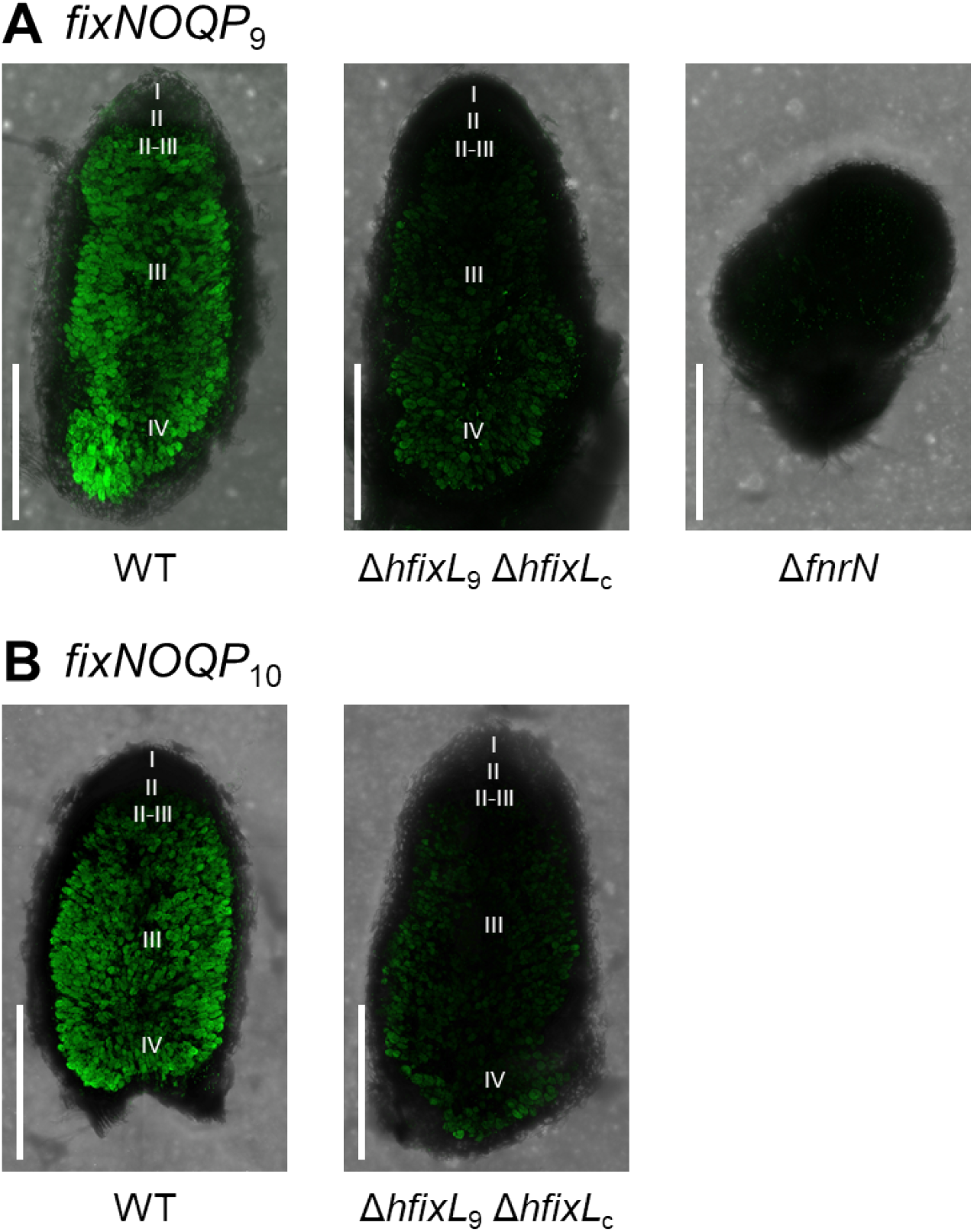
Spatial expression pattern of the *fixNOQP* operons in nodules infected with Rlv 3841 WT and mutants. (**A**) Expression of *fixNOQP*_9_ in strains of Rlv 3841. In nodules inoculated with WT, expression starts abruptly at the II-III interzone. In the double *hfixL* mutant, expression is reduced and begins gradually and deeper within the nodule, in zone III. Almost no expression is found in *fnrN* mutant nodules. (**B**) Expression of *fixNOQP*_10_ in WT Rlv 3841 and the double *hfixL* mutant followed a similar pattern as *fixNOQP*_9_. Expression begins at the II-III interzone. In the double *hfixL* mutant, expression begins deeper, in zone III of the nodule, and is reduced. Images within a set were captured and processed using identical parameters. Fluorescence intensity between the A and B image sets should not be compared as intensity was normalised. See Materials and Methods for microscopy technique details.

### 2.4. Integration of FnrN and hFixL improves the O_2_ response

To study the dynamics of an integrated cascade containing both hFixL and FnrN, we constructed an ordinary differential equation model of the system based on past literature (Supplementary 2). A map of the regulatory connections incorporated in the model is given in Supplementary 1, Figure S3. One homolog each of hFixL, FxkR, FixK, FnrN and FixNOQP is considered in the model. hFixL was assumed to become active near a headspace concentration of 1% O_2_ and FnrN near 0.01% O_2_, corresponding to dissolved O_2_ concentrations of ~12 μM and ~120 nM respectively at equilibrium. Past studies have suggested that hFixL binds O_2_ cooperatively, and that FnrN and FixK bind DNA as dimers [61,89–92]. Consequently all three processes were modelled using Hill functions with a Hill coefficient of 2 [93]. FixK and FnrN were assumed to have an identical induction effect on transcription when bound to the anaerobox motif. However, based on the critical role that FnrN but not FixK plays in regulation, FnrN was assumed to have a greater binding affinity for anaerobox motifs than FixK.

Our model reproduced the biphasic response of the integrated hFixL-FnrN cascade observed in WT Rlv3841 as O_2_ dropped from atmospheric (~21%) to near-anoxic concentrations (0.001 %) (Figure 8A-B). Expression of *fnrN* and *fixNOQP* first began at microaerobic conditions (10-1% O_2_) under the action of the hFixL-FxkR-FixK system. Subsequent activation of FnrN around 0.01% O_2_ led to a further increase in *fixNOQP* expression, mirroring our findings *in planta*. Expression of *fnrN* near 0.01% O_2_ initially increased due to auto-activation. Subsequently, as the O_2_ concentration continued to drop, *fnrN* expression decreased due to auto-repression. Thus, the model correctly predicts a generally homogenous level of *fnrN* expression throughout the nodule, and increased *fixNOQP* expression in the core of the nodule relative to the tip.

**Fig. 8.**
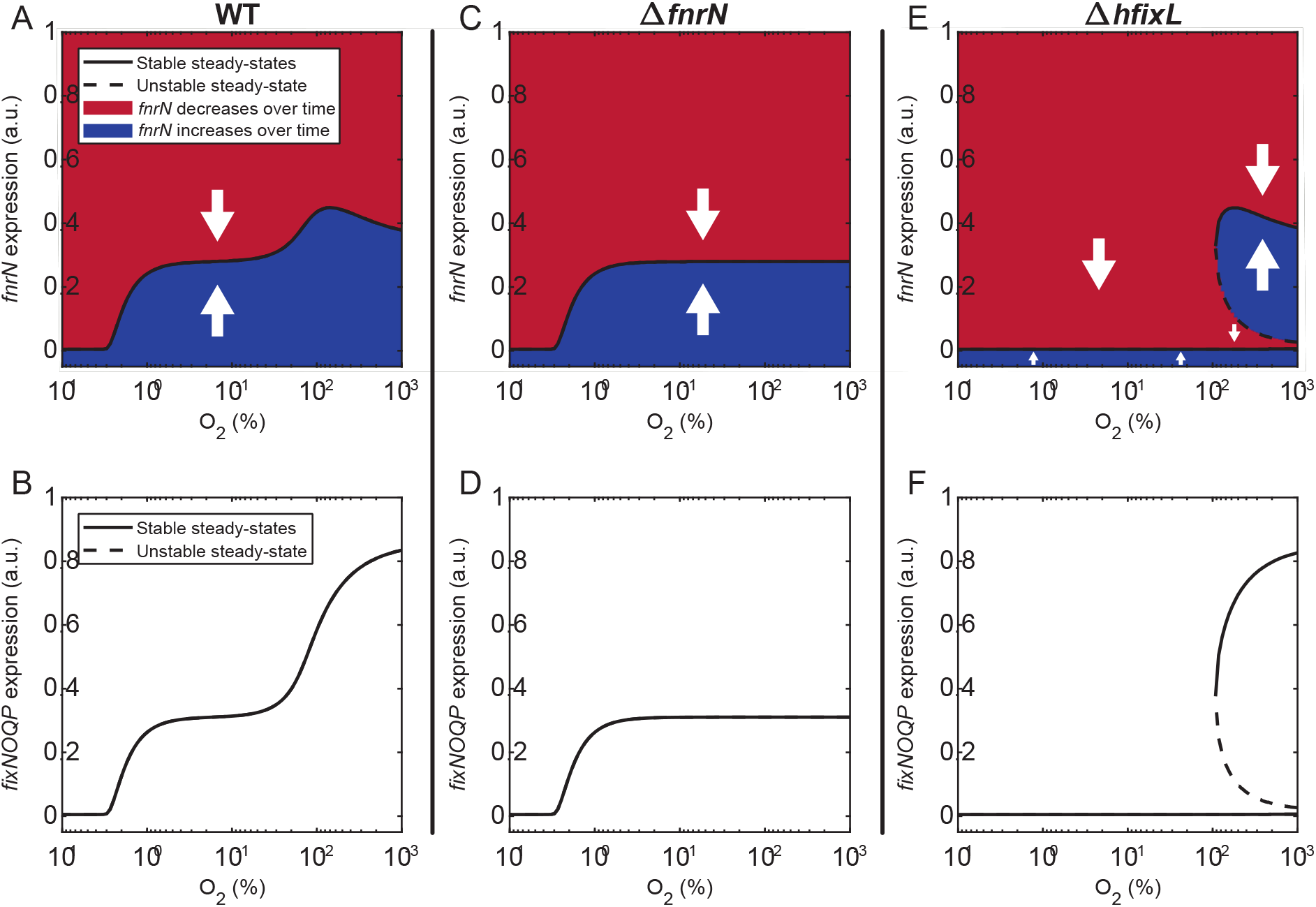
Modelling predicts the biphasic response of *fnrN* and *fixNOQP* expression controlled by a cascade integrating the hFixL and FnrN oxygen sensors. (**A, C, E**) Expression of *fnrN*. (**B, D, F**) Expression of *fixNOQP*. Black lines indicate stable expression steady-states, dashed lines indicate unstable steady states. In cells whose state is in a red shaded area, expression is expected to decrease; in a blue shaded area, it is expected to increase. (**A**) In a WT system comprising both sensors, expression of *fnrN* initially begins under microaerobic conditions. Expression increases slightly due to FnrN auto-activation as O_2_ concentration drops, but is stabilized by auto-repression. (**B**) Expression of *fixNOQP* begins under microaerobic conditions, then increases when FnrN becomes active as O_2_ drops further. (**C, D**) In the absence of FnrN, expression of the *fnrN* gene and *fixNOQP* operon begins under microaerobic conditions driven by the hFixL-FxkR-FixK pathway, but does not subsequently increase. (**E, F**) In cells where only the FnrN system is present, expression of the *fnrN* gene and *fixNOQP* operon does not occur until O_2_ concentration drops sufficiently for FnrN to be active. Once FnrN is active, the system exhibits bistability, with stable states of both near-zero and high levels of *fnrN* and *fixNOQP* expression. As the O_2_ concentration continues to drop, an increasing proportion of cells are expected to transition to the high expression state due to stochastic variations in expression.

In the absence of FnrN (Figure 8C-D), the model shows initial expression of *fnrN* and *fixNOQP* under microaerobic conditions due to the hFixL-FxkR-FixK pathway. There is however no further induction as O_2_ continues to drop. This agrees with our finding that some *fnrN* expression takes place in the Rlv3841 *fnrN* mutant, albeit at a reduced level relative to WT. The model also predicts a lower level of *fixNOQP* expression in the *fnrN* mutant, consistent with our confocal microscopy results for *fixNOQP*_9_ expression (Figure 7A).

In the absence of *hfixL* (Figure 8E-F), an important new behaviour of the system is predicted by our model. As expected, no induction of *fnrN* or *fixNOQP* takes place under microaerobic conditions, in line with our experimental findings. However, once O_2_ drops below 0.01%, our model suggests that the system may be bistable, with possible steady states at either high *fnrN* expression or near-zero expression. As the O_2_ concentration continues to decrease, the disturbance needed to move from minimal expression to the high expression steady state becomes smaller. Thus, as the bacteria experience increasingly anaerobic conditions in deeper parts of the nodule, the model predicts that an increasing proportion of cells will transition from near-zero to high *fnrN* expression due to stochastic variations in expression of the gene. This agrees with the gradual increase in *fixNOQP* expression observed in the *hfixL* double mutant in Rlv3841 (Figure 7A-B).

## 3. Discussion

O_2_ regulation is essential for rhizobia to establish a successful symbiosis with their legume partners. The model *Rhizobium* Rlv3841 employs three O_2_ sensors in symbiosis, hFixL, FnrN and NifA. In the present study, we examined this multi-sensor system through a combination of *in vivo, in vitro* and *in silico* approaches. The hFixL-FxkR-FixK system is active in the earliest stages, followed by FnrN as the bacteria enter deeper into the nodule. Both regulate genes required for symbiotic survival, such as *fixNOQP*. NifA is active at a later stage, in zone III of nodules, and regulates activation of core nitrogen fixation machinery. The hFixL-FxkR-FixK system is the most O_2_ tolerant of the three sensors, active in free-living bacteria under microaerobic conditions and *in planta* beginning in zone I of nodules. The FnrN protein is inactive under free-living microaerobic conditions and only becomes active from the II-III interzone onwards. FnrN is critical for expression of *fixNOQP* and nitrogen fixation activity. Indirectly, the hFixL-FxkR-FixK system also plays an important role by inducing *fnrN* expression under microaerobic conditions, priming it for auto-activation in the central nitrogen fixing zone. Our modelling results suggest the hFixL-FxkR-FixK system also prevents bistability in the low O_2_ response, thereby ensuring all cells commit to *fixNOQP* expression within the central nitrogen fixing zone. Thus hFixL-FxkR-FixK and FnrN act as a single regulation pathway which integrates both O_2_ sensors.

Rhizobia experience a drop in O_2_ concentration of at least three orders of magnitude as they transition from a free-living lifestyle in soil to terminally differentiated bacteroids in nodules. Like Rlv3841, it is common for rhizobia to employ multiple O_2_ sensors during this transition. These multiple sensors may be used to create redundancy, a feature often found in key regulatory pathways to improve their robustness. Elements of this redundancy are present in Rlv3841, including the multiple hFixL homologs and the overlap between their role and that of FnrN. However, our results also demonstrate that each sensor plays an important distinct role. Thus, integrating sensors into a single cascade in Rlv3841 also improves the responsiveness of regulation and allows the bacteria to respond appropriately across the entire range of O_2_ concentrations experienced during symbiosis. A similar dual-sensor arrangement has also previously been described in *Rhodopseudomonas palustris*, which combines a FixLJ-FixK cascade with the FnrN homolog AadR [94–96]. *R. palustris* is not symbiotic but is noted for its ability to grow under both aerobic and anaerobic conditions [97]. The combined pathway in *R. palustris* was shown to provide fine-tuned regulation for adapting to the large range of O_2_ concentrations it experiences.

The prevalence of multi-sensor O_2_ regulation systems in rhizobia may also have arisen in response to competitive fitness pressures. Legume plants can sanction rhizobia based on their nitrogen fixation activity [98–100]. The bacteria may also be selected based on the speed with which they are able to adapt to life inside nodules and begin productively fixing nitrogen. This would create pressure for strains to rapidly demonstrate their effectiveness to their legume host. By enabling more fine-tuned control, integrated multi-sensor O_2_ regulatory systems may speed up the symbiotic transition, providing a competitive advantage.

## 4. Materials and Methods

### Bacterial strains and growth conditions

*E. coli* strains were grown in liquid or solid Luria-Bertani (LB) medium[101] at 37 °C supplemented with appropriate antibiotics (μg mL^-1^): ampicillin 100, kanamycin 20, spectinomycin 50 and gentamicin 10. Rlv3841 strains were grown at 28 °C in Tryptone-Yeast (TY) extract[102] or Universal Minimal Salts (UMS)[103] with glucose and ammonium chloride at 10 mM each. Antibiotics for Rlv3841 were used at the following concentrations (μg mL^-1^): gentamicin 20, kanamycin 50, spectinomycin 100, streptomycin 500, tetracycline 2, neomycin 80 and nitrofurantoin 20.

### Cloning, colony PCRs and conjugations

All routine DNA analyses were done using standard protocols [101]. PCR reactions for cloning were carried out according to manufacturer’s instructions with Q5 High-Fidelity DNA Polymerase (New England Biolabs). Colony PCRs used OneTaq DNA Polymerase (NEB). Restriction enzymes (NEB) were used according to the manufacturer’s instructions. Sanger sequencing was carried out by Eurofins Genomics. Assemblies using BD In-Fusion cloning (Takara Bio) were performed according to the manufacturer’s instructions. Conjugations and transductions with bacteriophage RL38 were performed as previously described [104,105]. Tn7 integrations were performed according to the method described by Choi and colleagues [106,107].

### Mutant generation and complementation

#### Rlv3841 *hfixL*_c_ (RL1879) mutant, LMB403

A 1 Kb internal fragment of *hfixL*_c_ was PCR amplified from Rlv3841 with primers pr0988/0989, adding XbaI sites at the 5’ and 3 ends. This fragment was cloned into pK19mob digested with XbaI, using BD In-Fusion cloning, to produce plasmid pLMB441. Triparental filter conjugation of pLMB441 into WT Rlv3841 was then performed using kanamycin selection. Colonies were screened with colony PCR using primers pr0482 and pK19A, which bind upstream of *hfixL*_c_ and inside the integrated pK19 backbone respectively. This gave mutant strain LMB403.

#### Rlv3841 *hfixL*_9_ (pRL90020) mutant, LMB495

A 1 Kb region containing *hfixL*_9_ was PCR amplified from Rlv3841 with primers pr1270/1271. This region was subcloned into pJET1.2/blunt to produce plasmid pLMB581. Plasmid pLMB581 was then digested with XbaI/XhoI and the *hfixL*_9_ region cloned into pJQ200SK using BD In-Fusion, digested with the same enzymes, to give plasmid pLMB585. A spectinomycin resistance cassette was digested out of the pHP45ΩSpc plasmid with SmaI and cloned into pLMB585 at a unique StuI site blunted using the Klenow fragment to give plasmid pLMB590. Triparental filter conjugation of pLMB590 into WT Rlv3841 was then performed using spectinomycin selection. Colonies were screened with colony PCR using primers pr1272/1273. This gave mutant strain LMB495.

#### Rlv3841 double *hfixL*_c_ *hfixL*_9_ mutant, LMB496

An Rlv3841 mutant in both *hfixL* genes was generated by triparental filter conjugation of pLMB441 into strain LMB495, producing double mutant strain LMB496.

#### Rlv3841 *fnrN* (RL2818) mutant, LMB648

A 2.5 Kb region containing *fnrN* was PCR amplified from Rlv3841 with primers pr1381/1382. This fragment was digested with XbaI/XhoI and cloned using BD In-Fusion into pJQ200SK linearized with digestion by the same enzymes to make plasmid pLMB732. A tetracycline resistance cassette was then digested out of the pHP45ΩTc plasmid with EcoRI and cloned into pLMB585 at a unique MfeI site to give plasmid pLMB733. Triparental filter conjugation of pLMB733 into WT Rlv3841 was then performed using tetracycline selection. Colonies were screened with colony PCR using primers pr1432/1433. This gave mutant strain LMB648.

#### Rlv3841 triple *hfixL*_c_ *hfixL*_9_ *fnrN* mutant, LMB673

A triple Rlv3841 mutant, in both *hfixL* genes and the *fnrN* gene, was generated by transducing *fnrN*ΩTc from LMB648 into LMB496 to produce strain LMB673.

#### Rlv3841 *fxkR*_9_ (pRL90026) mutant, OPS1808

Two 1 Kb regions, one upstream and one downstream of *fxkR*_9_ were PCR amplified from Rlv3841 with primer pairs oxp2874/2875 and oxp2876/2877 respectively. These were cloned with BD In-Fusion into pK19mobSacB digested with PstI and EcoRI to produce plasmid pOPS1199. Triparental filter conjugation of pOPS1199 into WT Rlv3841 was then performed using kanamycin selection. Colonies were screened with colony PCR using primers oxp3155 and pK19A. Colonies with correct integration were subsequently subjected to sucrose selection to remove plasmid pK19mobSacB as previously described [108]. Colonies were then screened for loss of kanamycin resistance and using colony PCR with primers oxp3155/3156 to isolate mutant strain OPS1808.

#### Complemented Rlv3841 *fnrN* mutant (OPS2260)

The *fnrN* gene with its native promoter was amplified from Rlv3841 with primers oxp4115/4116 and cloned into BsaI-digested pOGG280 using BD In-Fusion. This plasmid was then genomically integrated with kanamycin selection and colonies screened with primers oxp2327/2328 and confirmed with sequencing. This produced strain OPS2260.

#### *In vitro* microaerobic induction measurement

Rlv3841 strains were first grown on slopes with appropriate antibiotics for three days. Cells were resuspended and washed three times by centrifugation at 5,000 RCF for 10 minutes. Washed cells were used to inoculate 10 mL liquid UMS cultures to OD_600_ 0.01 and grown overnight without antibiotics. Cultures were then diluted to OD 0.1 in 400 μL UMS each in a 24-well microtiter plate (4titude). A gas-permeable membrane (4titude) was applied to plates. Plates were then incubated in a FLUOstar Omega plate reader equipped with an Atmospheric Control Unit (both produced by BMG) to adjust O_2_ concentration to 1% and CO_2_ concentration to 0.1%. Readings were taken every 30 minutes and plates shaken at 700 rpm in double orbital mode between readings. Induction was measured at 18 hrs postinoculation, when all cultures had reached stationary phase.

#### Plant growth and acetylene reduction

*Pisum sativum* cv. Avola seeds were surface sterilized using 95% ethanol and 2% sodium hypochlorite before sowing. Plants were inoculated with 1 × 10^7^ cells of the appropriate rhizobial strain and grown in 1 L beakers filled with sterile medium-grade vermiculite and nitrogen-free nutrient solution as previously described in a growth room (16h light/8h dark) [109]. Harvesting was 21 days later and acetylene reduction rate was determined as previously described [110]. All nodules for each plant were counted and weighed; acetylene reduction rates were normalised by total nodule weight.

### Statistical analysis

All analyses were performed using GraphPad Prism 8 (GraphPad Software). Significant differences were determined by Student’s t-test or one-way ANOVA followed by Dunnett’s multiple comparisons post-hoc test correction. A p-value less than 0.05 was considered statistically significant.

### Confocal microscopy

Reporters were constructed by transcriptional fusion of promoters to an ORF of the sYFP2 fluorescent protein. Reporters were subsequently genomically integrated into Rlv3841 strains using the mini-Tn7 system [106]. Plants were inoculated with marked strains and grown as described above. After 21 days, nodules were picked and immersed in water then cut in half longitudinally. Images were taken with an LSM 880 confocal laser-scanning microscope equipped with the Axio Imager.Z2 (Zeiss), using the manufacturer’s ZEN Black software. A Plan-Apochromat 10×/0.45 M27 objective (Zeiss) was used. Excitation was at 514 nm with an Argon laser and emission measurements filtered to a range of 519-572 nm. Acquisitions were tile scans with 2×3 tiles per image. 31 Z-stack slices were taken at each slice, separated by a height of 10 μm. Images shown in this publication are maximum-intensity orthogonal projections produced with the ZEN Blue software (Zeiss).

## Acknowledgements

The authors would like to thank Dr Tim Haskett, Dr Carmen Sánchez-Cañizares and Prof Lee Sweetlove for their advice and critically reviewing the manuscript. They would also like to thank Dr Niloufer Irani for her help with confocal microscopy, and Dr Beatriz Jorrín for her help with the Tn7-based reporters used in this study.

## Supporting information captions

1. **S1 Text. Supporting figures**
2. **S2 Text. Modelling oxygen regulation in Rlv3841**
3. **S3 Text. Strains, plasmids and primers used in the study**

